# Functional and signaling characterization of the neutrophil FPR2 selective agonist Act-389949

**DOI:** 10.1101/571604

**Authors:** Simon Lind, Martina Sundqvist, Rikard Holmdahl, Claes Dahlgren, Huamei Forsman, Peter Olofsson

**Affiliations:** Department of Rheumatology and Inflammation Research, Institute of Medicine, University of Gothenburg, Sweden; Medical Inflammation Research, Department of Medical Biochemistry and Biophysics, Karolinska Institute, Stockholm, 17177 Sweden

**Author notes:** Correspondence to: Claes Dahlgren Department of Rheumatology and Inflammation Research, University of Gothenburg, Box 480, S-405 30 Göteborg, Sweden. These authors contributed equally to the work (Huamei Forsman, Peter Olofsson).

**Keywords:** neutrophils, formyl peptide receptors, receptor agonists, NADPH-oxidase, β-arrestin

## Abstract

Despite the steadily increased numbers of formyl peptide receptor (FPR) ligands identified over the years, few have been characterized in studies using animal disease models and even less have entered clinical trials in human subjects. A small-molecule compound, Act-389949, was however recently tested in a phase I clinical trial and found to be safe and well tolerated in healthy human subjects. The desired anti-inflammatory property of Act-389949 was proposed to be mediated through FPR2, one of the FPRs expressed in neutrophils, but no basic characterization was included in the study. To gain more insights into FPR2 recognition of this first-in-class compound for future utility of the agonist, we have in this study determined the receptor preference and down-stream signaling characteristics induced by Act-389949 in human blood neutrophils isolated from healthy donors. Our data demonstrate that Act-389949 is an agonist for FPR2 that triggers functional/signaling repertoires comparable to what has been earlier described for other FPR2 agonists, including neutrophil chemotaxis, granule mobilization and activation of the NADPH-oxidase. In fact, Act-389949 was found to be as potent as the prototype FPR2 peptide agonist WKYMVM and had the advantage of being resistant to oxidation by the MPO-H_2_O_2_-halide derived oxidants, as compared to the sensitive WKYMVM. The down-stream signals generated by Act-389949 include an FPR2-dependent and Gαq-independent transient rise in intracellular Ca^2+^ and recruitment of β-arrestin. In summary, our data show that Act-389949 serves as an excellent tool-compound for further dissection of FPR2-regulated activities *in vitro* and *in vivo*. Potent and stable FPR ligands such as Act-389949 may therefore be used to develop the next generation of FPR signaling regulating anti-inflammatory therapeutics.

## Introduction

Neutrophil granulocytes, the most abundant leukocytes in human blood, are key players in regulating and fine-tuning inflammatory reactions [1, 2]. The functional repertoires of these cells are triggered and mediated through cell surface receptors of which the formyl peptide receptors (FPRs), belonging to the family of G-protein coupled receptors (GPCRs) are the most extensively studied [3, 4]. Human neutrophils express two FPRs, FPR1 and FPR2, which both are high affinity receptors for formylated peptides, a molecular pattern hallmark of proteins/peptides encoded for by bacterial as well as mitochondrial genes [3, 4]. This type of danger signals are recognized by the FPRs during microbial infections, inflammation, cancer, and tissue injury [5, 6]. In addition to formyl peptides, FPRs recognize also a number of non-formyl peptides and other molecules belonging to different chemical classes including small compounds and lipopeptides [7-9]. Among the numerous FPR specific/selective ligands reported, many transduce pro-inflammatory activities in neutrophils by mediating directional cell migration (chemotaxis), secretion of granule contents and, activation of the superoxide generating electron transporting NADPH-oxidase [10]. The immunoregulatory effects and impact of FPR mediated activation and release of reactive oxygen species (ROS) has for decades solely been considered as pro-inflammatory and harmful. Yet, the complexity of the regulatory FPR down-stream signaling and secretion of ROS has during recent years been reevaluated. This is in part based on that a deficiency of one of the subunits of the NADPH oxidase complex, leading to an inability to generate ROS, results in a life-threatening immunocompromised condition termed chronic granulomatous disease (CGD) [11]. In addition to suffering from repeated severe infections, CGD patients also have an increased risk of developing autoimmune/inflammatory diseases [12-14], suggesting an inflammation-dampening effect of ROS. We have previously through positional cloning of genetic polymorphism found that *Ncf1* (encoding for the p47^phox^ subunit of the NADPH oxidase complex) is disease-associated and linked to a lower capacity to produce ROS [15-17], which is of importance for disease severity in animal models of arthritis (15, 16), psoriasis [18], colitis [19], and lupus [20]. Interestingly, polymorphism of *Ncf1* plays a similar role in human autoimmune diseases [21]. Accordingly, our accumulated research proposes a regulatory role of ROS produced by the NADPH-oxidase in many cellular processes [22, 23]. One of the major mechanisms utilized for activation of NADPH oxidase produced ROS is via activation of FPRs [3], which could result in a ROS mediated regulation of inflammatory responses. In addition to activation of NADPH oxidase, it has been shown that FPRs mediate also anti-inflammatory/resolving activities when interacting with peptides cleaved off from the N-terminus of the calcium regulated protein annexin I [24, 25] as well as with arachidonic acid derived lipid agonists [26], suggesting that FPRs can transduce distinct ligand-directed responses. In line with this, we and others have demonstrated that GPCRs, including FPR2, selectively can couple to the β-arrestin or G-protein signaling pathways and mediate biased signaling responses [27-29]. The functional selective response induced by the lipopeptide/pepducin F2Pal10, that triggers a rise in intracellular calcium ([Ca^2+^]i) and activates the superoxide generating NADPH-oxidase but is unable to recruit β-arrestin and induce chemotaxis [28], nicely illustrates the biased signaling concept. Hence, due to the complex signaling triggered by FPR agonists a detailed *in vitro* characterization of FPR agonists is critical for understanding the effects of FPR modulating drugs and a prerequisite for further preclinical and clinical development of FPR agonists as drug candidates.

Despite the large effort to identify small molecules that potently and selectively activate FPR2, the success rate has been very low. However, there are some examples of identified and characterized potent FPR agonists. Actually, one of the first small compound identified as an FPR2 specific agonist (Quin-C1) was shown to possess biased signaling properties as it induced a transient rise in [Ca^2+^]i but no oxidase activity. This compound was also shown to dampen inflammation in a mouse lung injury model [30-32]. Another commonly used small compound, Cmp43, originally identified through an FPR2 based screening process [33], turned out to be a dual agonist for both FPR1 and FPR2 [34] with FPR1 as the preferred receptor in primary human neutrophils [35]. The majority of the identified FPR agonists have so far only been characterized in *in vitro* experiments and to our knowledge only one compound, Act-389949, has entered clinical trial. This small compound drug candidate developed by Actelion Pharmaceuticals Ltd, was described to be a potent and selective FPR2 agonist and it was introduced in a phase I clinical trial comprising healthy human subjects [36]. The compound was shown to be safe and well tolerated but the drug-potential was regarded to be hampered by the fact that it desensitized FPR2, and the transient response was interpreted to be pro-inflammatory rather than anti-inflammatory [36]. A detailed *in vitro* characterization of this presumed FPR2 agonist, critical for further clinical development of stable FPR2 agonists as inflammatory modulatory therapeutics was, however, not included in the published study. Hence, in this study we have performed a detailed *in vitro* characterization of the small molecule Act-389949 and determined the receptor preference and signaling profile in human neutrophils. Our results not only confirm the FPR2 preference over FPR1 but also provide mechanistic insights into FPR2-mediated signaling events and cellular responses by Act-389949. To our knowledge, and as demonstrated by data provided in this study, Act-389949 is one of the most potent FPR2 agonist belonging to the small compound class known today. Thus, Act-389949 could serve as a valuable tool-compound for further mechanistic studies and FPR2-based development of clinically active compounds.

## MATERIAL AND METHODS

### Ethics Statement

This study, conducted at the Sahlgrenska Academy in Sweden, includes peripheral blood and blood from buffy coats obtained from the blood bank at Sahlgrenska University Hospital, Gothenburg, Sweden. According to the Swedish legislation section code 4§ 3p SFS 2003:460 (Lag om etikprövning av forskning som avser människor), no ethical approval was needed since the blood samples were provided anonymously and could not be traced back to a specific donor.

### Chemicals and reagents

Dextran and Ficoll-Paque were from Pharmacia (Uppsala, Sweden) and Fura-2-AM was from Life Technologies Europe (Stockholm, Sweden). RPMI 1640 culture medium without phenol red was purchased from PAA Laboratories GmbH (Pasching, Austria). Isoluminol, tumour necrosis factor α (TNFα), N-formyl-Met-Leu-Phe (fMLF), cetyltrimethylammonium bromide (CTAB), o-Phenylenediamine (OPD), EGTA, dimethyl sulfoxide (DMSO), platelet activating factor (PAF), bovine serum albumin (BSA) and Latrunculin A were obtained from Sigma-Aldrich (St. Louis, MO, USA). Horse radish peroxidase (HRP) and superoxide dismutase (SOD) were purchased from Boehringer-Mannheim (Mannheim, Germany). IL8 was from R&D systems (Minneapolis, MN, USA) and the FPR2 specific hexapeptide Trp-Tyr-Met-Val-Met-NH2 (WKYMVM) was synthesized and purified by HPLC by Alta Bioscience (University of Birmingham, Birmingham, United Kingdom). The FPR2 specific antagonist PBP_10_ and agonist F2Pal_10_ were synthesized by CASLO Laboratory (Lyngby, Denmark) and the FPR1 specific inhibitor (an inverse agonist) cyclosporin H was kindly provided by Novartis Pharma (Basel, Switzerland). The phycoerythrin (PE) conjugated anti-CD62L antibody and allophycocyanin (APC) conjugated anti-CD11b antibody were purchased from Becton Dickinson Biosciences (Sparks, MD, USA). The Gαq inhibitor YM-254890 was purchased from Wako Chemicals (Neuss, Germany) and the phenylacetamide compound (S)-2-(4-chlorophenyl)-3,3-dimethyl-N-(5-phenylthiazol-2-yl)butanamide (Cmp58) was obtained from Calbiochem-Merck Millipore (Billerica, MA, USA). The Act-389949 compound (N-(2-{[4-(1,1-difluoroethyl)-1,3-oxazol-2-yl]methyl}-2H-1,2,3-triazol-4-yl)-2-methyl-5-(3-methylphenyl)-1,3-oxazole-4-carboxamide), synthesized by Ramidus AB (Lund, Sweden) as described in WO2010/143116 (p69; [36, 37], was a generous gift from ProNoxis AB (Lund, Sweden). Myeloperoxidase (MPO) was a kind gift from Inge Olsson (Lund, Sweden). The peptides/receptor antagonists were dissolved in DMSO to a concentration of 10^−2^ M and stored at -80°C until use. Further dilutions were made in Krebs-Ringer phosphate buffer containing glucose (10 mM), Ca^2+^ (1 mM), and Mg^2+^ (1.5 mM) (KRG; pH 7.3).

### Isolation of human neutrophils

Neutrophil granulocytes were isolated from peripheral blood or buffy coats obtained from healthy adults [38]. After dextran sedimentation at 1 x *g*, hypotonic lysis of the remaining erythrocytes, and centrifugation on a Ficoll-Paque gradient, the neutrophils were washed and re-suspended (1 x 10^7^/ml) in KRG. The cells were stored on melting ice until used. In some of the experiments, neutrophils were primed with TNFα (10 ng/ml, 37°C, 20 minutes) before utilized.

### Calcium mobilization

Neutrophils at a density of 5 x 10^7^ cells/ml in KRG without Ca^2+^ supplemented with 0.1% BSA were loaded with 5 µM Fura-2-AM for 30 minutes in the dark at room temperature. The cells were then diluted 1:1 in RPMI 1640 culture medium without phenol red and centrifuged. Finally, the cells were washed once with KRG and re-suspended in the same buffer to a density of 2 x 10^7^/ml. Calcium measurements were carried out in a PerkinElmer fluorescence spectrophotometer (LC50, Perkin Elmer, Waltham, MA, USA), with excitation wavelengths of 340 nm and 380 nm, an emission wavelength of 509 nm, and slit widths of 5 nm and 10 nm, respectively. The transient rise in intracellular calcium is presented as the ratio of fluorescence intensities (340 nm/380 nm) detected [39].

### Neutrophil NADPH-oxidase activity

Neutrophil superoxide anion production was determined using an isoluminol-enhanced chemiluminescence (CL) system [40, 41]. The CL activity was measured in a six-channel Biolumat LB 9505 (Berthold Co, Wildbad, Germany) using disposable 4-ml polypropylene tubes with a 1-ml reaction mixture. Tubes containing isoluminol (2 x 10^−5^ M), HRP (2 units/ml), and neutrophils (10^5^/ml) were equilibrated for five minutes at 37°C, after which 0.1 ml of stimuli was added and the superoxide production, measured as light emission and expressed in Mega counts per minute (Mcpm), was recorded continuously over time.

By a direct comparison of the superoxide dismutase (SOD) reduction of cytochrome C to CL, 7.2 x 10^7^ counts per minute were found to correspond to a production of 1 nmol of superoxide as measured with the cytochrome C reduction technique (a millimolar extinction coefficient for cytochrome C of 21.1 was used; further details about the CL technique is given in [40, 42].

### Treatment of agonists with MPO-H_2_O_2_

Peptide agonist WKYMVM (10^−6^ M) or Act-389949 (10^−6^ M) diluted in KRG was incubated with MPO (1 μg/ml) at ambient temperature for five min before the addition of H_2_O_2_ (final concentration, 10 μM), and the samples were incubated for another 10 min to allow peptide oxidation at ambient temperature. The remaining activity of the agonists after MPO-H_2_O_2_-halide oxidation was determined through their potential to trigger the ROS release in neutrophils.

### Chemotaxis assay

Neutrophil migration was determined by a Boyden chamber technique using 96-well microplate chemotaxis chambers containing polycarbonate filters with 3 μm pores (Chemo-Tx; Neuro Probe Inc., Gaithersburg, MD, USA) according to manufacturer’s instructions. In short, Act-389949 and WKYMVM, diluted in KRG buffer supplemented with 0.3% BSA, were added to wells in the lower chamber. Cell suspensions (30 μl) containing neutrophils (2 x 10^6^/ml, isolated from peripheral blood) were placed on top of the filter and allowed to migrate for 90 min at 37°C. The cell migration to the bottom well was visualized under microscope and for quantitative analysis the content of MPO was assessed in the lysates (cells in lower chamber treated with 2% BSA and 2% CTAB for 60 minutes, at room temperature) by addition of OPD and hydrogen peroxide. Triplicate samples for each stimulus was performed and all values were subtracted by the negative control (spontaneous migration of neutrophils towards buffer). The data is presented as percent migration as compared to the positive control (neutrophils added directly to the bottom chamber, i.e., 100% migration).

### Cell surface receptor exposure of CD62L and CD11b

Neutrophils (5 x 10^6^/ml) were incubated with various concentrations of Act-389949 or fMLF as the positive control at 37°C for 10 minutes. Thereafter, the cells were co-incubated with a PE conjugated anti-CD62L antibody (1:40 dilution) and an APC conjugated anti-CD11b (1:20 dilution) for 45 minutes on ice. The cells were then washed, re-suspended after which the cell surface exposure of CD62L and CD11b was determined on an Accuri C6 flow cytometer (Becton Dickinson Biosciences, Sparks, MD, USA) and analyzed by FlowJo software version 10.3 (Tree star Inc., Ashland, Oregon, USA).

### β-arrestin recruitment assay

The ability of agonists in promoting FPRs to recruit β-arrestin was evaluated in the PathHunter® eXpress CHO-K1 FPR1 or FPR2 cells from DiscoverX (Fremont, CA, USA) which co-express ProLink tagged FPR1 or FPR2 and an enzyme acceptor tagged β-arrestin so that β-arrestin binding can be measured via enzyme fragment complementation as increased β-galactosidase activity. The assay was performed according to manufacturer’s instructions and as previously described [28, 43]. In brief, cells were diluted in a Cell Plating Reagent, seeded in tissue culture treated 96-well plates (10.000 cells/well) and incubated at 37°C, 5% CO_2_ for 20 hours. The cells were then incubated with agonists (90 minutes, 37°C), followed by addition of detection solution (60 minutes, room temperature) and measurement of chemiluminescence on a CLARIOstar plate reader (BMG Labtech, Ortenberg, Germany).

### Data analysis

Data analysis was performed using GraphPad Prism 8.0 (Graphpad Software, San Diego, CA, USA). Curve fitting was performed by non-linear regression using the sigmoidal dose-response equation (variable-slope). Statistical analysis was performed on raw data values using a repeated measurement one-way ANOVA (Figures 6A-B) or an ordinary one-way ANOVA (Figure 4A) followed by Dunnett’s multiple comparison or, a paired Student’s *t*-test (Insets Figures 6C, 7A-C). Statistically significant differences are indicated by \**p* < 0.05, \*\**p* < 0.01, \*\*\**p* < 0.001, \*\*\*\**p* < 0.0001 and non-statistically significance difference is indicated by ns.

**Fig 4.**
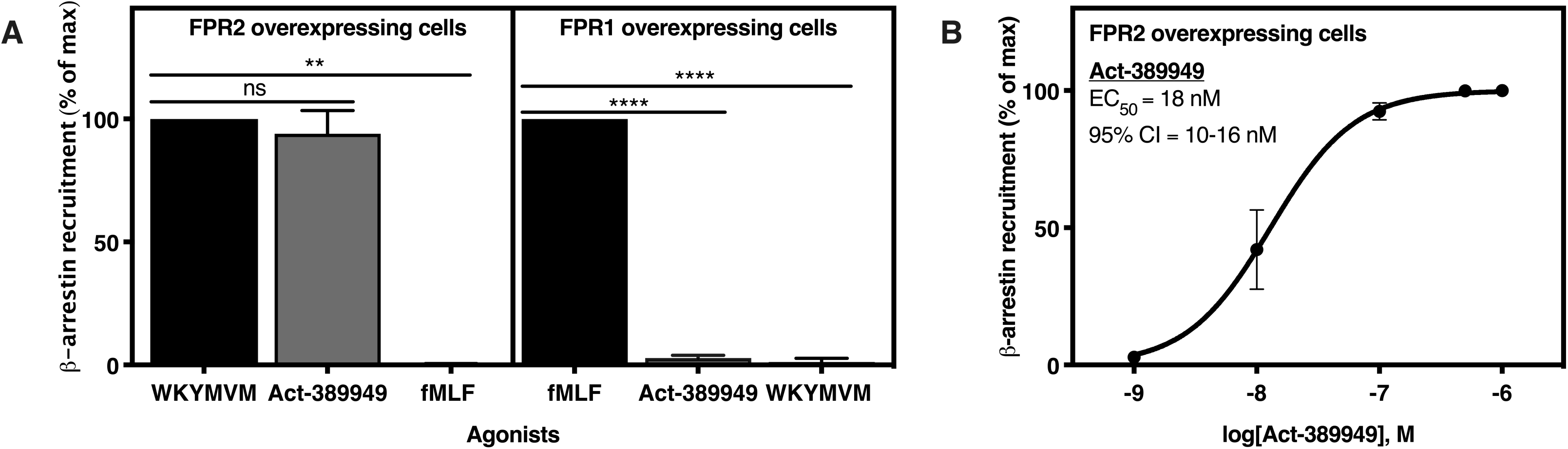
Act-389949 recruits *β*-arrestin in FPR2 overexpressing cells. FPR-mediated β-arrestin recruitment was determined in PathHunter® CHO cells. **A.** Response in FPR2 (left) and FPR1 (right) overexpressing CHO cells when stimulated with 100 nM of either WKYMVM, Act-389949 or fMLF. The data are presented as percent of the response induced by saturating concentrations of WKYMVM (100 nM) for FPR2 overexpressing cells and fMLF (100 nM) for FPR1 overexpressing cells; mean±SD, n=3). **B.**The responses induced in FPR2 overexpressing CHO cells when stimulated with different concentrations of Act-389949 were determined and used for calculation of the EC_50_-value, including the 95% confidence interval (CI), mean±SD (n=3).

**Fig 6.**
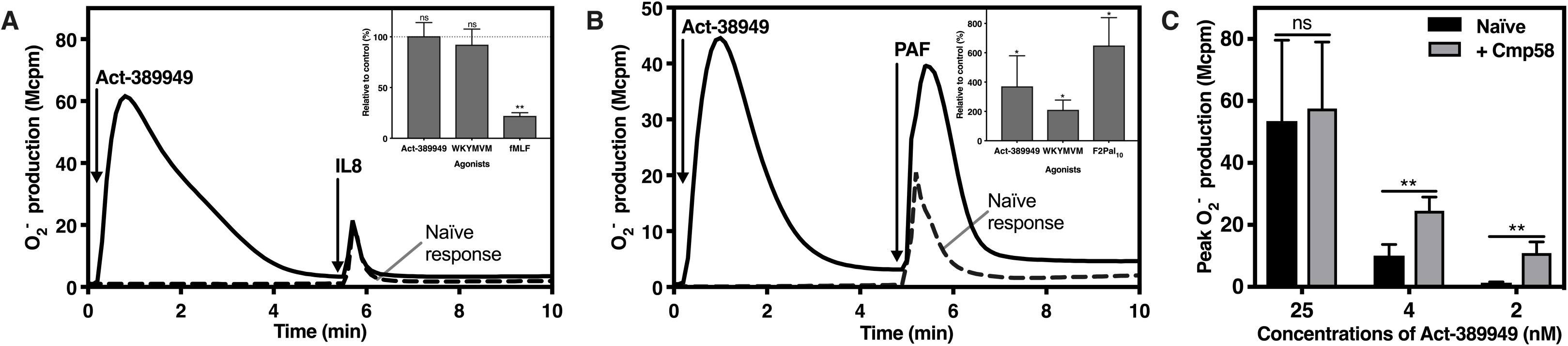
Act-389949 and NADPH-oxidase activity by receptor cross-talk mechanisms. The NADPH-oxidase activity induced in neutrophils was determined. Abscissa, Time (min); ordinate, O_2_^−^ production, arbitrary Mcpm units). **A-B.** The NADPH-oxidase responses induced by IL8 and PAF (broken lines) were compared with the responses induced when neutrophils were activated in sequence with Act-389949 (100 nM) and then with IL8 (100 ng/ml; arrow to the right in **A**) or PAF (100 nM; arrow to the right in **B**): The NADPH-oxidase mediated superoxide anion production was determined, and one representative experiment of at least three independent experiments is shown for each stimulation. Abscissa, Time (min); ordinate, O_2_^−^ production, arbitrary Mcpm units). **Insets (A):** The IL8 response induced in naïve neutrophils relative (percent) to the IL8 response induced in neutrophils pre-stimulated with 100 nM of either Act-389949, WKYMVM or fMLF (mean±SD, n=3). The dashed line indicates 100%. **Insets (B):** The PAF-induced response in naïve neutrophils relative (percent) to the PAF response induced neutrophils pre-stimulated with Act-389949 (100 nM), WKYMVM (100 nM) or F2Pal_10_ (500 nM) (mean±SD, n=3). The NADPH-oxidase activity was determined in neutrophils pre-incubated (5 min) with or without the allosteric FFA2R modulator Cmp58 (1 µM), when activated with different concentrations of Act-389949. The bar graphs represent the peak NADPH-oxidase activity induced by Act-389949 in naïve and Cmp58-modulated neutrophils, respectively (mean±SD; n=3).

## RESULTS

### Act-389949 triggers an FPR2-dependent and Gα*q independent rise of intracellular Ca*^*2+*^

The small compound agonist Act-389949, presumed to be a selective FPR2 activating molecule (Fig 1, [36]), was synthesized and its ability to induce a rise in the cytosolic concentration of free calcium ions ([Ca^2+^]_i_) in human neutrophils was determined. Act-389949 induced a transient rise in [Ca^2+^]_i_ in neutrophils (Fig 2A) with a concentration in the low nM range and the time course of the response very similar to that induced by the earlier described receptor selective peptide agonists fMLF (specific for FPR1) and WKYMVM (specific for FPR2) (Fig 2B). Earlier studies have shown that the rise in [Ca^2+^]_i_ induced by FPR agonists is achieved through the PLC-PIP_2_-IP_3_ signaling route in which the generated IP_3_ triggers a release of Ca^2+^ from intracellular stores [39]. This signaling route is largely independent of the extracellular concentration of Ca^2+^ and rely in neutrophils on receptor-coupling to a heterotrimeric G-protein that belongs either to the Gαi or the Gαq subfamily [44]. The independence of extracellular Ca^2+^ was illustrated by the fact that addition of EGTA (to chelate extracellular Ca^2+^) had no substantially effect of the rise in [Ca^2+^]_i_ induced by Act-389949 (Fig 2C), strongly suggesting that the activity is dependent on a release of Ca^2+^ from intracellular stores. Further, the selective Gαq inhibitor YM-254890 had no effect on the Act-389949 induced rise in [Ca^2+^]_i_ (Fig 2D; the inhibitory effect on the PAF/PAFR response, which exclusively involves Gαq [44], was included for comparison; Fig 2E), implying that the classical FPR signaling route mediated by the βγ heterodimer of a Gαi containing G-protein is triggered by Act-389949. In order to determine the precise receptor preference of Act-389949, the inhibitory effects of the receptor selective antagonists cyclosporin H (specific for FPR1 [45, 46]) and PBP_10_ (specific for FPR2 [47, 48]) were determined. Our data clearly show that FPR2 was the preferred receptor for Act-389949, as the Act-389949 induced [Ca^2+^]_i_ response was completely inhibited by PBP_10_ (Fig 2F, middle curve) but unaffected by cyclosporin H (Fig 2F, lower curve).

**Fig 1.**
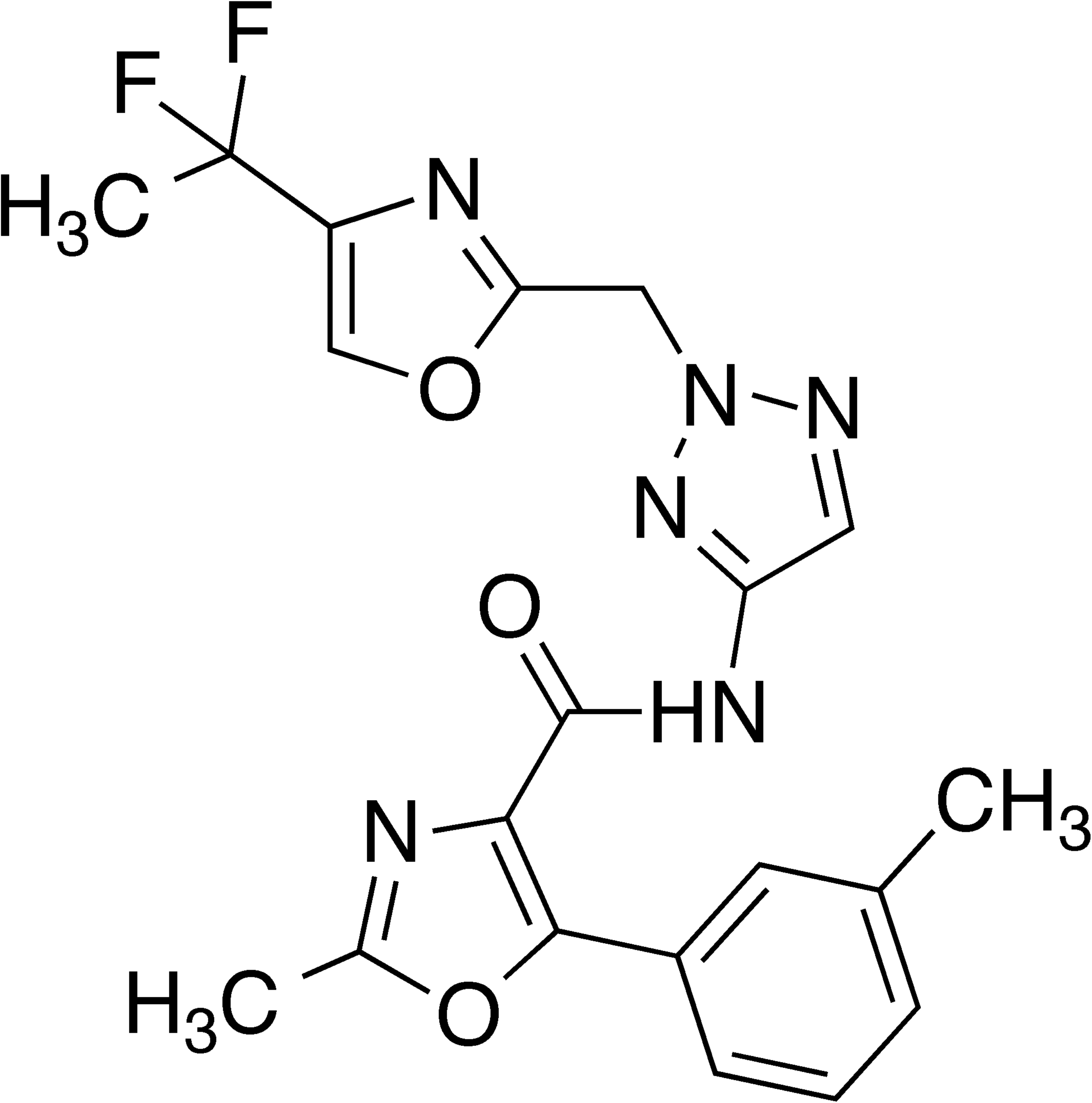
Chemical structure of Act-389949. The presumed FPR2 agonist Act-389949 described in a phase 1 clinical trial in healthy human subjects was synthesized. The calculated physicochemical properties of Act-389949 were: hydrogen bond acceptors = 7; hydrogen bond donors =1; rotatable bonds =7; topographic polar surface area =111.87; molecular weight = 428.14.

**Fig 2.**
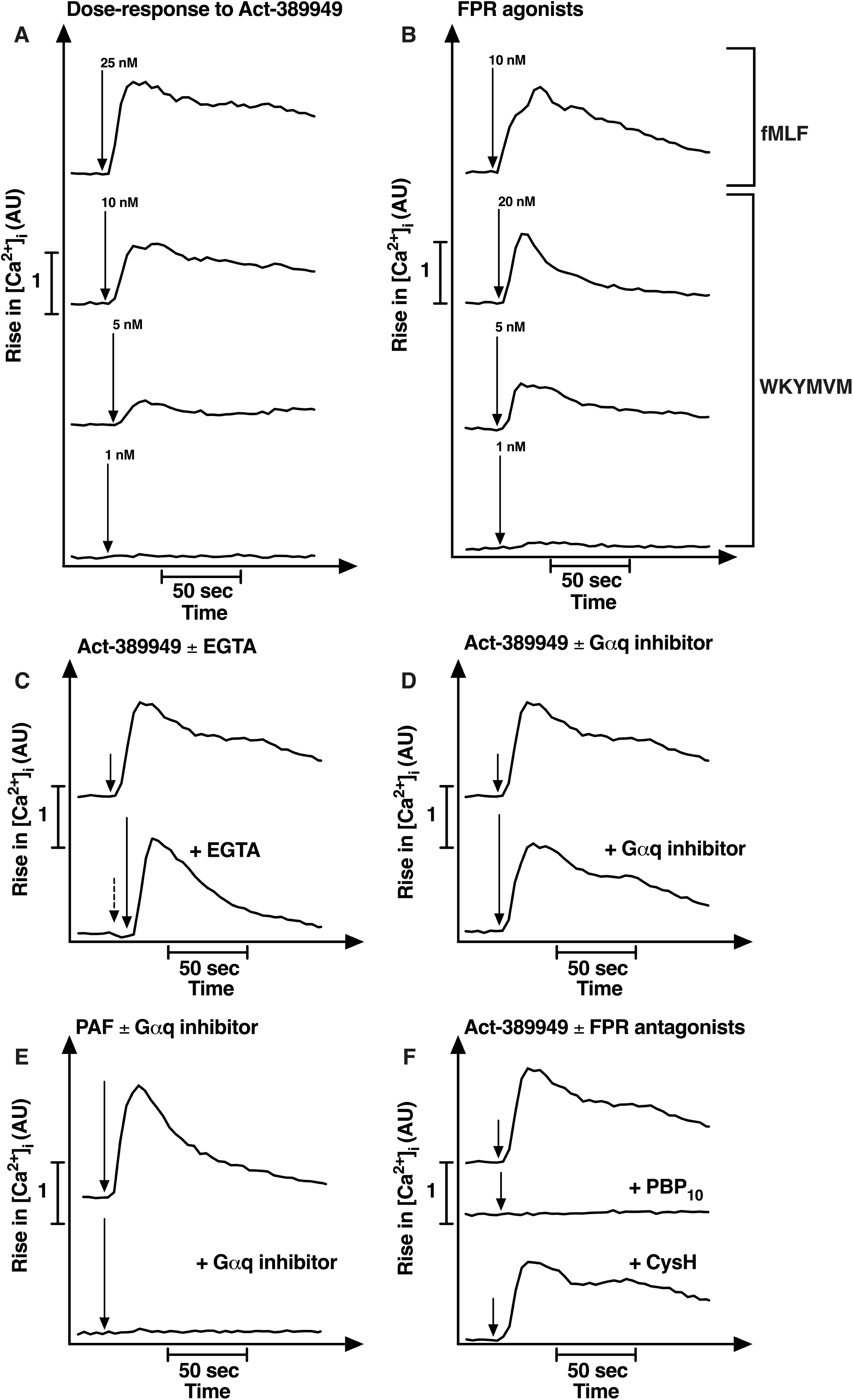
The transient rise in intracellular Ca^2+^ induced by Act-389949 is not inhibited by the Ca^2+^ chelator EGTA or the Gαq inhibitor YM-254890. Neutrophils were loaded with Fura-2 and stimulated with different agonists added as indicated by arrows. Abscissa, time of study (sec); ordinate, increase in intracellular Ca^2+^ ([Ca^2+^]_i_) given as the change in the ratio between Fura-2 fluorescence at 340 and 380 nm (AU, arbitrary units). **A**-**B**. The transient rise of [Ca^2+^]_i_ was followed in **A**) neutrophils stimulated with Act-389949 (different concentrations;1 nM-25 nM), **B**) the FPR1 agonist fMLF (upper panel; 10 nM) and the FPR2 agonist WKYMVM (lower panels; different concentrations;1 nM-20 nM). **C.** Neutrophils were stimulated with Act-389949 (10 nM) in the absence (upper curve) and presence of EGTA (2.5 mM; lower curve) added 5 sec before Act-389949 (indicated by the broken arrow) and the transient rise of [Ca^2+^]_i_ was followed. **D**-**E.** Neutrophils were stimulated with **D**) Act-389949 (10 nM) or **E**) PAF (1 nM) in the absence (upper curve) and presence of YM-254890 (a selective Gαq inhibitor; 200 nM, added 5 minutes before the agonist; lower curve) and the transient rise of intracellular Ca^2+^ was followed. **F.** The transient rise of [Ca^2+^]_i_ was followed in neutrophils stimulated with Act-389949 (10 nM) in the absence (upper curve) and presence of the FPR2 antagonist PBP_10_ (1 µM, added 5 minutes before the agonist; middle curve) or the FPR1 antagonist cyclosporin H (1 µM, added 5 minutes before the agonist; lower curve). (**A-F**) Representative traces from one of at least 3 independent experiments are shown.

### Act-389949 triggers an FPR2-dependent release of superoxide anions from human neutrophils

To study the FPR2-medidated neutrophil functional response triggered by Act-389949, we next investigated the ability of Act-389949 to induce release of NADPH-oxidase generated superoxide anions, as compared to the earlier characterized FPR1 agonist fMLF and, the FPR2 agonist WKYMVM [3, 4]. The three molecules induced very similar neutrophil respiratory burst responses, characterized by a very short lag phase followed by a rapid increase of superoxide release and a peak of activity reached after around one minute (Fig 3A). Of note, when a 100 nM concentration of each agonists was used, the magnitude of the responses induced by the three agonists were comparable (Fig 3A), indicating that Act-389949 is a full agonist. The respiratory burst induced by Act-389949 was concentration dependent with an EC_50_ value of around 10 nM (Fig 3B). In accordance with the profile of other FPR agonists, the response induced by Act-389949 was substantially increased in TNFα primed neutrophils (Fig 3C). The inhibitory profile with receptor specific antagonists corroborated the [Ca^2+^]_i_ (Fig 2F) by showing that FPR2 was the preferred receptor for Act-389949 (inhibition by PBP_10_ but not by cyclosporin H, Fig 3D; the inhibitory profile of WKYMVM and fMLF are added for comparison). Further, neutrophils activated with Act-389949 were not only desensitized in their response to a second dose of Act-389949, but also non-responsive to the FPR2 agonist WKYMVM but fully responsive to the FPR1 agonist fMLF (Fig 3E). In agreement with the [Ca^2+^]_i_ data obtained with the Gαq inhibitor (Fig 2D), the neutrophil NADPH-oxidase response to the FPR2 agonists WKYMVM, Act-389949 or the FPR2 biased lipopeptide (pepducin) F2Pal_10_ [28] was not affected by the Gαq inhibitor YM-254890 (Fig 3F). Taken together, these data demonstrate that Act-389949 is a potent and full FPR2 agonist that activates the neutrophil NADPH-oxidase independent of Gαq.

**Fig 3.**
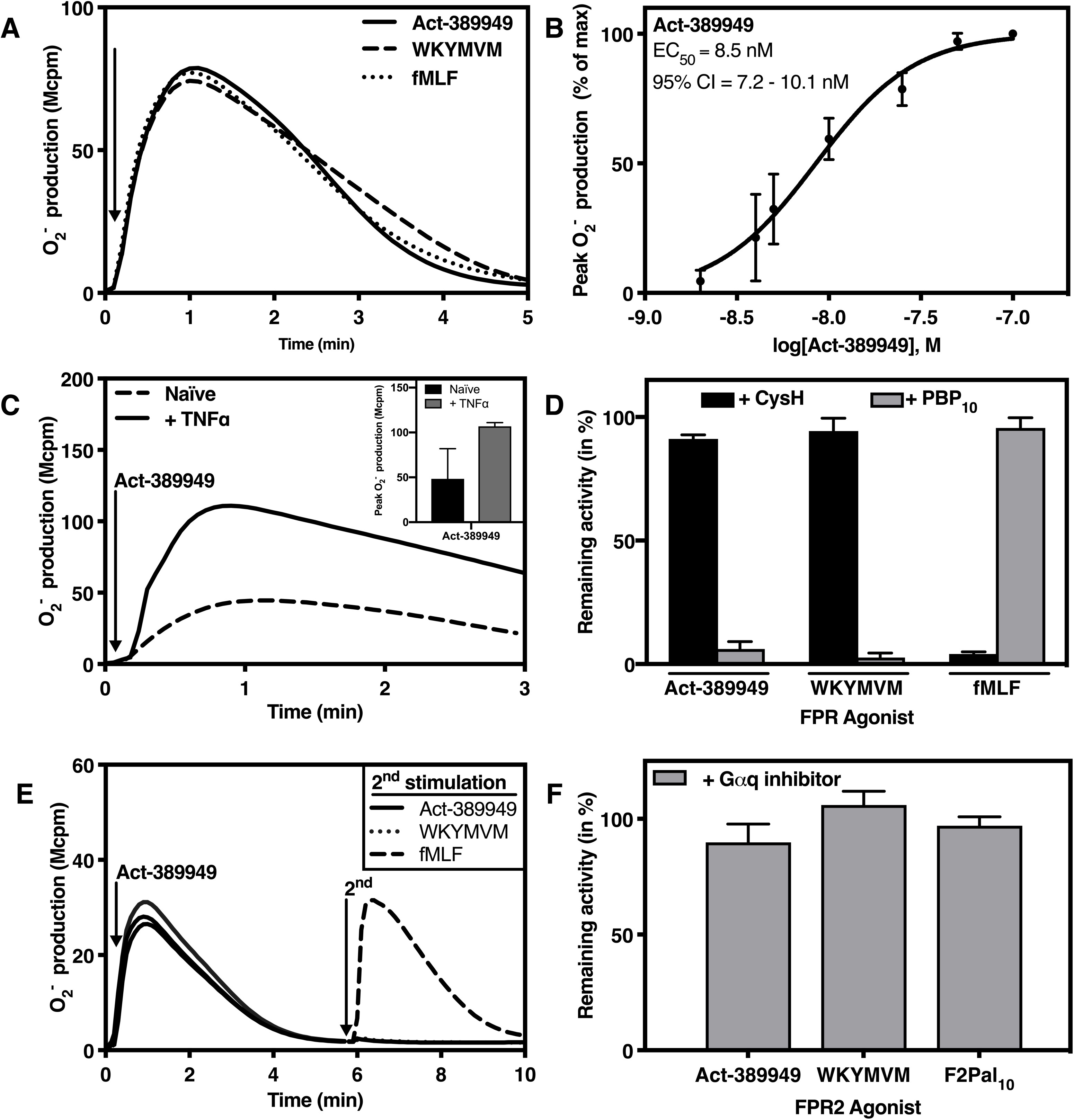
Activation of the neutrophil NADPH-oxidase by Act-309949. Neutrophils were pre-incubated at 37°C for five minutes (min) before stimulation (indicated by arrows) and the NADPH-oxidase mediated superoxide anion (O_2_^−^) production was determined Abscissa, Time (min); ordinate, O_2_^−^ production, arbitrary Mcpm units). **A.** Neutrophils were stimulated with 100 nM of either Act-389949, fMLF or WKYMVM. **B.** Neutrophils were stimulated with different concentrations of Act-389949 and the EC_50_-value, including the 95% confidence interval (CI), was determined based on the peak O_2_^−^-response. **C.** Naïve and TNFα primed neutrophils were challenged with Act-389949 (100 nM). **Inset:** Comparison between the peak O_2_^−^-responses released from naïve and TNFα primed cells. **D.** Comparison between the peak O_2_^−^-responses released by neutrophils in the absence or presence of either cyclosporin H or PBP_10_ (1 µM respectively) before stimulation with 100 nM of either Act-389949, fMLF or WKYMVM. The data are presented as percent remaining NADPH-oxidase activity in the presence of antagonists as compared to the responses from control cells. **E.** Neutrophils were first stimulated with Act-389949 (100 nM, arrow to the left) and then further challenged a second time (arrow to the right marked 2^nd^) with 100 nM of either Act-389949, WKYMVM or fMLF. **F.** Comparison between the peak O_2_^−^ responses released by neutrophils, treated with and without the Gαq inhibitor YM-254890 (200 nM, added during pre-incubation) before stimulation with Act-389949 (100 nM), WKYMVM (100 nM) or F2Pal_10_ (500 nM). The data are presented as percent remaining NADPH-oxidase activity in the presence of YM-254890, compared to the responses from control cells. The data show (**A, C** and **E**) one representative trace of at least three independent experiments for each stimulation, (**B**) mean±SD; n=3 and (**C inset, D** and **F**) mean±SD; n=3.

### Act-389949 promotes FPR2 but not FPR1 to recruit β-arrestin

Receptor recruitment of cytosolic β-arrestin, induced by agonist binding, is a common feature for many GPCRs and this process is suggested to be of importance not only for subsequent receptor desensitization and internalization, but also for the ability of the activated receptor to initiate β-arrestin-mediated novel signaling pathways [49]. Accordingly, we recently showed that the pepducin F2Pal_10_ is a biased FPR2 agonist that lacks ability to recruit β-arrestin and to trigger chemotaxis [28]. Similar to the FPR2 peptide agonist WKYMVM but, in contrast to F2Pal_10_ and the FPR1 selective agonist fMLF, Act-389949 triggered β-arrestin recruitment in cells overexpressing FPR2 (Fig 4A). In agreement with the above receptor preference profile, and in accordance to WKYMVM, but opposite to fMLF, Act-389949 was unable to induce β-arrestin recruitment in FPR1 overexpressing cells determined with concentrations up to 500 nM (Fig 4A). The FPR2-mediated β-arrestin recruitment induced by Act-389949 was concentration dependent with an EC_50_ value of around 20 nM (Fig 4B).

### Act-389949 is a chemoattractant and secretagogue for human neutrophils

Chemotaxis is a functional response induced by many, but not all FPR agonists, evident from the fact that the agonist WKYMVM but not F2Pal_10_ attracts neutrophils [28]. Activation of neutrophils, due to migration from blood to tissue or due to presence of inflammatory mediators, is in most instances associated with a reorganization of membrane surface receptors, e.g., CD62L (L-selectin), which is proteolytically shed from the surface upon activation or upregulation of granule-stored membrane proteins e.g. CD11b (complement receptor 3). This activation can be mimicked *in vitro* by incubation neutrophils with e.g., and TNFα and FPR agonists [50].

Act-389949 triggered neutrophil migration causing a typical bell-shaped dose-response curve, comparable to the conventional FPR2 agonists WKYMVM (Fig 5A-B). In addition, and similar to the prototype FPR2 agonist WKYMVM, Act-389949 concentration dependently triggered shedding of CD62L (Fig 5C-D) and induced degranulation resulting in increased surface exposure of CD11b (Fig 5E-F). Taken together, these data show that Act-389949 triggers not only a transient rise in [Ca^2+^]_i_, an activation of the neutrophil NADPH-oxidase but also chemotaxis and granule mobilization in neutrophils.

**Fig 5.**
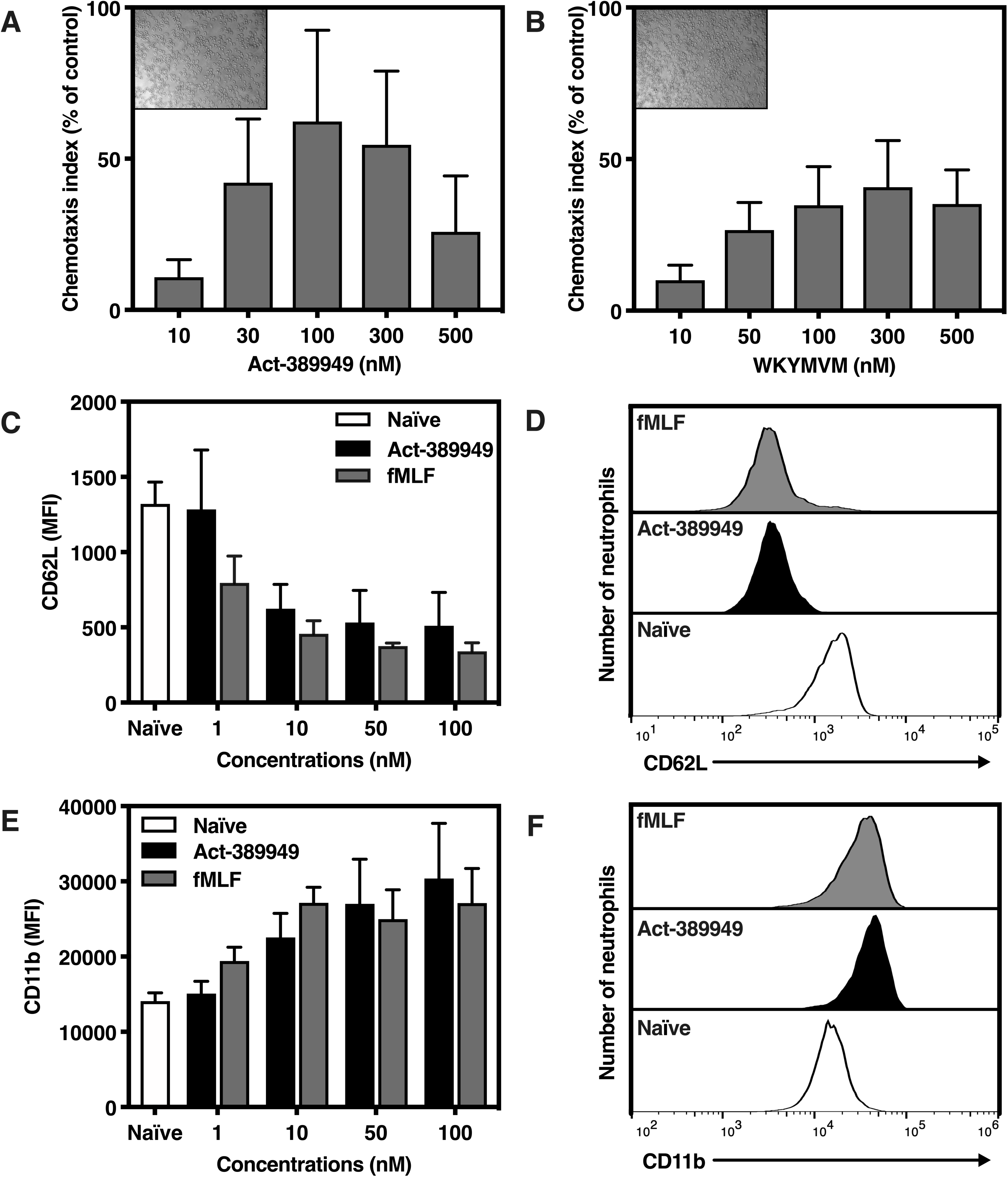
Act-389949 induces both neutrophil migration and degranulation. **A**-**B**. Neutrophils added to in the upper compartment (above the filter) were allowed to migrate towards different concentrations of Act-389949 (**A**) and WKYMVM (**B**) added to the lower compartment. The cells were allowed to migrate for 90 minutes and the number of cells in the lower compartment (below the filter) was determined by analyzing the amount of MPO. The bar graph represents the fraction of neutrophils that migrated through the filter (mean±SEM, n=3). I**nsets:** Representative microscopic images of neutrophils recovered in the lower compartment following migration towards 100 nM of either **A**) Act-389949 or **B**) WKYMVM. **C-F**. Act-389949 induced surface CD62L shedding and granule mobilization determined by the increase in surface expression of CD11b. Neutrophils were incubated for 10 min at 37°C in the absence and presence of Act-389949 or fMLF in concentrations ranging from 1-100 nM, before staining with anti-CD62L (**C**-**D**) or anti-CD11b (**E**-**F**) antibodies and analysis by flow cytometry. (**C** and **E**) The bar graphs show the mean fluorescence intensity (MFI) (mean±SD, n=3). (**D** and **F**) Representative histograms from one of the three donors.

### Act-389949 co-operates with other GPCR agonists and modulates the neutrophil NADPH-oxidase activity through receptor cross-talk mechanisms

There is a defined hierarchy between different neutrophil GPCRs [51], illustrated by the fact that neutrophils challenged with an FPR agonists are non-responsive to a second agonist that binds to the same receptor (homologous desensitization; see Fig 3). But, FPR1-desensitized neutrophils are also non-responsive to IL8, that binds CXCR1/2, a non FPR1 related GPCR (heterologous desensitization; Fig 6A inset and [3, 52]). No such heterologous desensitization was induced by Act-389949 on the IL8 response (Fig 6A); Act-389949 stimulated cells were responsive to a second stimulation with IL8 which agrees with the results obtained with the FPR2 peptide agonist WKYMVM (Fig 6A inset).

The neutrophil response can also be modulated through receptor-cross-talk mechanisms [44, 53, 54], a prominent example being that the agonist-occupied PAF receptors induce signals that induce reactivation of FPR2-desensitized neutrophils [44, 53]. In line with this, the PAF induced response was substantially increased in Act-389949 desensitized neutrophils compared to the response induced by PAF in naïve cells (Fig 6B). These results are in line with the known cross-talk pattern when using other FPR2 agonists, but the magnitude differs; the response induced by PAF in neutrophils desensitized by F2Pal_10_ was more pronounced than that induced in Act-389949 and WKYMVM desensitized neutrophils (Fig 6B inset).

To further explore Act-389949 mediated GPCR cross-talk in human neutrophils, we determined the Act-389949 induced neutrophil response in the presence of an FFA2R (free fatty acid receptor 2) specific allosteric modulator [54]. This allosteric modulator (Cmp58) has been shown to prime neutrophils for the responses induced by low concentrations of FPR agonists, a priming achieved through a novel receptor cross-talk mechanism [54, 55]. In accordance with published data, the NADPH-oxidase activity induced by low nanomolar concentrations of Act-389949 in neutrophils pre-incubated with the allosteric FFA2R modulator was also augmented compared to the response in naïve neutrophils (Fig 6C). In summary, Act-389949 bound FPR2 is able to cross-talk with other GPCRs in human neutrophils, as demonstrated by that Act-389949 amplifies the PAF response and that an FFA2R allosteric modulator primes the Act-389949 response.

### Regulation of the Act-389949-induced NADPH-oxidase response by the actin cytoskeleton

The NADPH-oxidase response mediated by many GPCRs including the FPRs and the ATP receptor P2Y_2_R is augmented in neutrophils treated with Cytochalasin B or Latrunculin A, drugs that disrupts the actin cytoskeleton [56, 57]. In addition, desensitized FPRs but not P2Y_2_R, are reactivated to produce NADPH-oxidase derived superoxide anions upon addition of a cytoskeleton-disrupting agent, revealing an actin-dependent desensitization/reactivation mechanism for FPRs [3]. The neutrophil response induced by Act-389949, similar to other FPR2 agonists such as WKYMVM and F2Pal_10_, was both increased and prolonged in the presence of Latrunculin A (Fig 7A and inset). Furthermore, the Act-389949 desensitized neutrophils were also reactivated by Latrunculin A (Fig 7B), in a comparable magnitude as WKYMVM and F2Pal_10_ (Fig 7B inset), suggesting that similar to other FPR2 agonists, desensitization of Act-389949 bound FPR2s also relies on an intact actin cytoskeleton.

**Fig 7.**
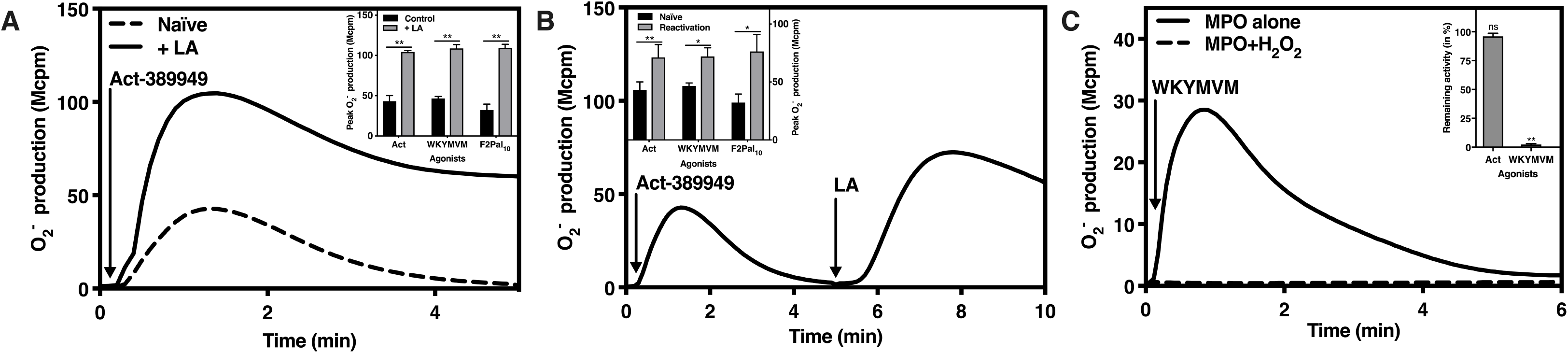
Modulation of the Act-389949 induced NADPH-oxidase response by Latrunculin A and the MPO/H_2_O_2_ system. The NADPH-oxidase activity induced in neutrophils was determined Abscissa, Time (min); ordinate, O_2_^−^ production, arbitrary Mcpm units). **A.** Naïve neutrophils and neutrophils incubated (5 min) with the actin cytoskeleton-disrupting drug Latrunculin A (LA; 25 ng/mL) were activated with Act-389949 (100 nM). One representative experiment of at least three independent experiments is shown. **Inset:** The peak NADPH-oxidase activities induced in naïve and LA treated neutrophils by Act-389949 (Act; 100 nM), or the FPR2 agonists WKYMVM (100 nM) and F2Pal_10_ (500 nM), respectively, were determine (mean±SD, n=3). **B.**Neutrophils activated by Act-389949 (100 nM, addition indicated by the arrow to the left) were reactivated with LA (25 ng/mL, addition indicated by arrow to the right). One representative experiment of at least three independent experiments is shown. **Inset:** The peak NADPH-oxidase activity induced by Act-389949 (Act; 100 nM), or the FPR2 agonists WKYMVM (100 nM) and F2Pal_10_ (500 nM), respectively, in naïve neutrophils were determined and compared with the peak NADPH-oxidase activity induced by reactivation with LA (mean±SD, n=3). **C** Neutrophils were stimulated with untreated or MPO/H_2_O_2_ treated WKYMVM (100 nM). One representative experiment of at least three independent experiments is shown. **Inset:** The peak NADPH-oxidase activity induced in neutrophils activated by MPO/H_2_O_2_ treated Act-389949 (Act; 100 nM) and WKYMVM (100 nM), respectively, compared to the activity induced by the untreated agonists (remaining activity in percent; mean±SD, n=3).

### Act-389949 is resistant to oxidation by the MPO-ROS-halide system

When the neutrophil superoxide/hydrogen peroxide (H_2_O_2_), generated by the NADPH-oxidase, are processed by the azurophil granule protein MPO, highly reactive oxidants are formed, and these constitute not only powerful weapons used to kill invading bacteria but can also inhibit/inactivate certain receptor agonists. Accordingly, we have previously shown that peptide agonists are rapidly inactivated by such oxidants [58, 59]. However, in comparison to WKYMVM, Act-389949 was stable when incubated together with MPO and H_2_O_2_, as determined by the ability of Act-389949 to activate neutrophils to produce superoxide after such a treatment (Fig 7C inset). That is, no inactivation was seen for Act-389949, whereas WKYMVM was inactivated, as no neutrophil NADPH-oxidase activation was achieved with this FPR2 agonist after incubation with MPO together with H_2_O_2_ (Fig 7C). In summary, the small compound FPR2 agonist Act-389949 but not the peptide FPR2 agonist WKYMVM is resistant to oxidation by the MPO-H_2_O_2_-halide system.

## Discussion

FPR2 is expressed by neutrophils and triggers chemotaxis, granule mobilization as well as activation of the NADPH-oxidase to produce reactive oxygen species. We have found that the small molecule Act-389949 is a selective FPR2 agonist that triggers a typical FPR2 functional repertoire in human neutrophils. A large number of publications have highlighted the immunomodulatory role of FPR2 by having both pro- and anti-inflammatory activities [1, 2, 60]. Thus, agonists as well as antagonists that target FPR2 may have therapeutic potential [61-64]. Accordingly, numerous FPR2 ligands have been identified and characterized through different approaches, but not until very recently, the agonist termed Act-389949 stated to be FPR2 specific, was introduced into a phase I clinical trial [36]. Act-389949 was found to be safe and well tolerated, and based on the fact that the number of receptors exposed on the cell surface of blood neutrophils/monocytes was reduced by Act-389949, rapid internalization of FPR2 and functional selective receptor desensitization were assumed [36]. However, no functional neutrophil responses or receptor preference data were included in the published study [36]. Consequently, to characterize the basic FPR activation properties of Act-389949 we have determined the receptor preference of Act-389949 and characterized the signaling and activation profile of the molecule in primary human neutrophils. We could confirm that Act-389949 is a potent and selective FPR2 agonist. This conclusion was based on data obtained with i) cells desensitized with WKYMVM, a well-known FPR2 agonist, ii) effects of the receptor specific inhibitor PBP_10_ (an FPR2 antagonist) that completely inhibited the neutrophil response whereas the FPR1 specific antagonist cyclosporine H was without effect and, iii) the ability of Act-389949 to recruit β-arrestin in cells overexpressing FPR2 but not FPR1. At the signaling level in neutrophils, the receptor-induced activities triggered by Act-389949 were very similar to those described earlier for conventional FPR2 specific agonists [3, 4]. A very early signal down-stream of activated FPRs is generated by the PLC-PIP_2_-IP_3_ signaling pathway and results in a transient rise in [Ca^2+^]_i_. This is achieved primarily through an IP_3_ induced emptying of the intracellular Ca^2+^ stores [65] and, accordingly, this pathway was triggered also by Act-389949. Furthermore, the fact that the [Ca^2+^]_i_ signaling was insensitive to a Gαq selective inhibitor suggests that the βγ subunits of a Gαi containing G-protein transduce the signal [44].

Over the last years it has become clear that biased agonism or functional selectivity is an essential concept in receptor signaling. The basis for receptor-specific biased agonists is that they selectively trigger one signaling pathway over another and induce a restricted/directed functional response. The molecular basis for this phenomenon has been suggested to be due to the different receptor sub-conformations induced by the agonist, however the precise mechanism is still not known. In line with the concept of biased FPR agonism, it has been suggested that activation of FPR2 transduce different down-stream signals when activated by conventional agonists and pro-resolving lipid ligands – the latter suggested to be biased ligands towards recruitment of β-arrestin, however, the effects of the lipid ligands are most likely mediated through a not yet identified receptor different from FPR2 [3]. Among the numerous small compound agonists described, one of the first identified non-peptide FPR2 agonist termed Quin-C1 [30] displays a biased property by lacking the ability to trigger superoxide release, while being able to induce both chemotaxis and degranulation in human neutrophils [30]. It is clear that also the FPR2-activating pepducin F2Pal_10_, is a biased FPR2 agonist and that the effects of F2Pal_10_ on neutrophil function differ in several aspects when compared to those mediated by conventional FPR2-specific peptide agonist [28]. Since the specificity or dual agonism of FPR1/2 molecules is essential for down-stream signaling and potential role in regulation of an immunological response, we aimed to characterize the concept of biased agonism for Act-389949. Our data demonstrate that the FPR2 signaling triggered by Act-389949 differs from that of Quin-C1 and F2Pal_10,_ yet was very similar (or even identical) to that induced by WKYMVM, a conventional peptide agonist for FPR2. These data strongly imply that Act-389949 should be regarded as a balanced FPR2 agonist that activates both G-protein and β-arrestin dependent signals.

Our data also show that FPR2 activating compounds including Act-389949 trigger a primed NADPH-oxidase response in neutrophils treated with TNFα. These cells are primed, not only in response to Act-389949, but also to various chemoattractants recognized by FPR1 and FPR2 [3, 4]. The primed response may be a regulatory mechanism, granting a cellular response (such as release of oxygen radicals) only at sites where it is functional and necessary, e.g., in inflamed or infected areas. Thus, the local ROS production *in vivo* upon FPR2 agonist stimulation may differ from *in vitro* and to a higher extent depend on the surrounding environment that determines the naïve or primed cellular state of the affected cells. Although the molecular mechanism(s) responsible for induction of the primed state is unclear, the exposure of new receptors to the plasma membrane due to degranulation is an attractive model as the molecular basis for the augmented response. This model is also in line with our earlier studies demonstrating that intracellular organelles are mobilized to the cell surface during TNFα priming [50], resulting in an increased exposure of various receptors including CD11b, FPR1, and FPR2 that are stored in the same intracellular organelles [3]. The molecular background to the TNFα induced primed response to Act-309949 may, thus, be the result of an increased exposure of FPR2 based on our data showing an increased surface exposure of CD11b upon Act-309949 stimulation.

Compared to peptide agonists, small compounds have certain advantages in therapeutic use due to their stability against degradation. In addition to proteolytic cleavage, we and others have earlier shown that some peptide FPR agonists can trigger their own inactivation due to oxidation by the MPO-hydrogen peroxide system [58, 66]. Our data show that Act-389949 is resistant to oxidation contrasting the conventional peptide agonist WKYMVM that transforms into a dormant agonist by the MPO and H_2_O_2_ system.

In summary, we provide certain molecular insights into Act-389949 induced FPR2 signaling and activation in human neutrophils. Our study demonstrates several important pro-inflammatory features of Act-389949, the first-in-class FPR2 agonist tested for its therapeutically potential in a clinical trial. Due to the complexity of the not yet completely understood biology and desensitization mechanism of FPR signaling, it is essential to understand and characterize functional signaling induced by compounds such as Act-389949. However, it is clear that selective and potent FPR2 agonists, like Act-389949, could be the starting material for an optimization process that together with different structure analogs may serve as valuable research tools for mechanistic studies both *in vivo* and *in vitro* where a stable agonist is required.

## Acknowledgements

This work was supported by the Swedish Research Council, The Swedish government under the ALF-agreement, the IngaBritt and Arne Lundberg research foundation, Eurostars project rosa GBS-E 10082, and Swedish Governmental Agency for Innovation Systems (project number 2016-01010) and Magnus Bergwall’s foundation (project number 2018-02579). We thank Malin Hultqvist Hopkins and Caroline Alkner at Redoxis AB (Lund, Sweden) for technical assistance.

## Authorship contributions

SL and MS performed the experiments and analyzed the data. RH, PO, CD and HF designed the study, and CD, HF wrote the manuscript which all authors commented on before approving the final version.

